# Basolateral localization of MMP14 drives apicobasal polarity change during EMT independently of its catalytic activity

**DOI:** 10.1101/402180

**Authors:** Cyril Andrieu, Audrey Montigny, Dominique Alfandari, Eric Theveneau

## Abstract

The transmembrane Matrix Metalloproteinase MMP14/MT1-MMP is known to promote cell migration by cleavage of the extracellular matrix. To initiate migration, epithelial cells need to gain mesenchymal attributes. They reduce cell-cell junctions and apicobasal polarity and gain migratory capabilities. This process is named epithelial-mesenchymal transition (EMT). MMP14’s implication in EMT is still ill-defined. We used chick neural crest (NC) cells as a model to explore the function of MMP14 in physiological EMT. Our results show that MMP14 is expressed by chick NC cells. However, it is its subcellular localization, rather than its expression, that correlates with EMT. MMP14 is first apical and switches to basolateral domains during EMT. Loss of function and rescue experiments show that MMP14 is involved in EMT independently of its catalytic activity. It lies downstream of pro-EMT genes and upstream of cell polarity. We found that basolateral localization of MMP14 is required and sufficient to induce polarity change in NC cells and neuroepithelial cells, respectively. These effects on polarity occur without impact on cell-cell adhesion or the extracellular matrix. Overall, our data points to a new function of MMP14 in EMT that will need to be further explored in other systems such as cancer cells.

## Introduction

MMP14 is a membrane-bound metalloproteinase (a.k.a Membrane Type1 Metalloproteinase, MT1-MMP) commonly upregulated in human cancer (Castro-Castro et al., 2016). It is primarily seen as a highly pro-invasive protease due to its cooperation with MMP2. Indeed, MMP14 is a direct activator of MMP2 (Sato et al., 1994) which can cleave collagen IV, a key component of the basal membranes of epithelial tissues (Halfter et al., 2015). Numerous studies have demonstrated the ability of MMP14 to promote invasion via its catalytic activity (Cathcart et al., 2016; Chen et al., 2016; Macpherson et al., 2014). However, its known functions are not restricted to proteolysis of matrix components in the context of cell migration. Studies in mammary gland development showed that MMP14 was dispensable for cell migration and branching but was required for deposition of white fat cells (Feinberg et al., 2016). In muscular dystrophy, MMP14 regulates the fibroblasts-adipocytes fate switch downstream of the Hedgehog pathway (Kopinke et al., 2017). Further, the cytoplasmic tail of MMP14 contains binding sites for regulators of the small GTPases Rac1 (Gonzalo et al., 2010a) and RhoA (Hoshino et al., 2009). Furthermore, MMP14 can interact directly with integrins (Gonzalo et al., 2010b). Besides these extracellular activities, MMP14 can be imported into the nucleus and displays transcriptional activities (Shimizu-Hirota et al., 2012). Most of these non-canonical functions have yet to be fully explored. In epithelial cancer, dissemination is initiated by an epithelial-mesenchymal transition (EMT). This complex process involves changes in adhesion, polarity and local matrix remodeling. MMPs are seen as late players in this process mostly for their ability to degrade the matrix to promote invasion/migration. However, their actual direct involvement in EMT remains poorly studied in vivo.

Here we used the chick neural crest (NC) cells as a model of physiological EMT. We found that MMP14 expression does not specifically associate with EMT or migration but that its subcellular localization changes during EMT. We demonstrate that the complete extracellular domain of MMP14, including its catalytic domain, is required for successful EMT but that catalytic activity itself is dispensable. MMP14 loss of function experiments show that MMP14 apical localization is not needed for the maintenance of a normal apicobasal polarity. However, basolateral localization of MMP14 is required and sufficient to impair apicobasal polarity in NC cells and neuroepithelial cells, respectively.

## Results

### MMP14 expression is not specifically associated with EMT and migration but its subcellular localization changes during EMT

In order to study the expression of MMP14, we cloned three partial cDNA of the coding sequence of chick MMP14 to generate probes for in situ hybridization. All probes gave similar results, representative images obtained with one of them are shown on Figure 1A-E. Wholemount staining indicate that MMP14 is expressed in the entire prospective brain (Fig. 1A). On transversal sections, MMP14 is found in the neural tube (Fig. 1B), including the neural crest domain (Fig. 1B’), and the notochord (Fig. 1B, arrow). Expression is maintained during EMT of NC cells (Fig. 1C) and is detected in early migratory cells (Fig. 1C’, arrowheads). At later stages (HH13), MMP14 is expressed in the brain, the eyes (e), the otic vesicle (ov) and the migratory NC cells (Fig. 1D, arrows). Longitudinal sections confirmed the presence of MMP14 mRNA in the migratory NC streams from rhombomeres (r) 1/2, 4 and 6 (Fig. 1E, arrows).

**Figure 1.**
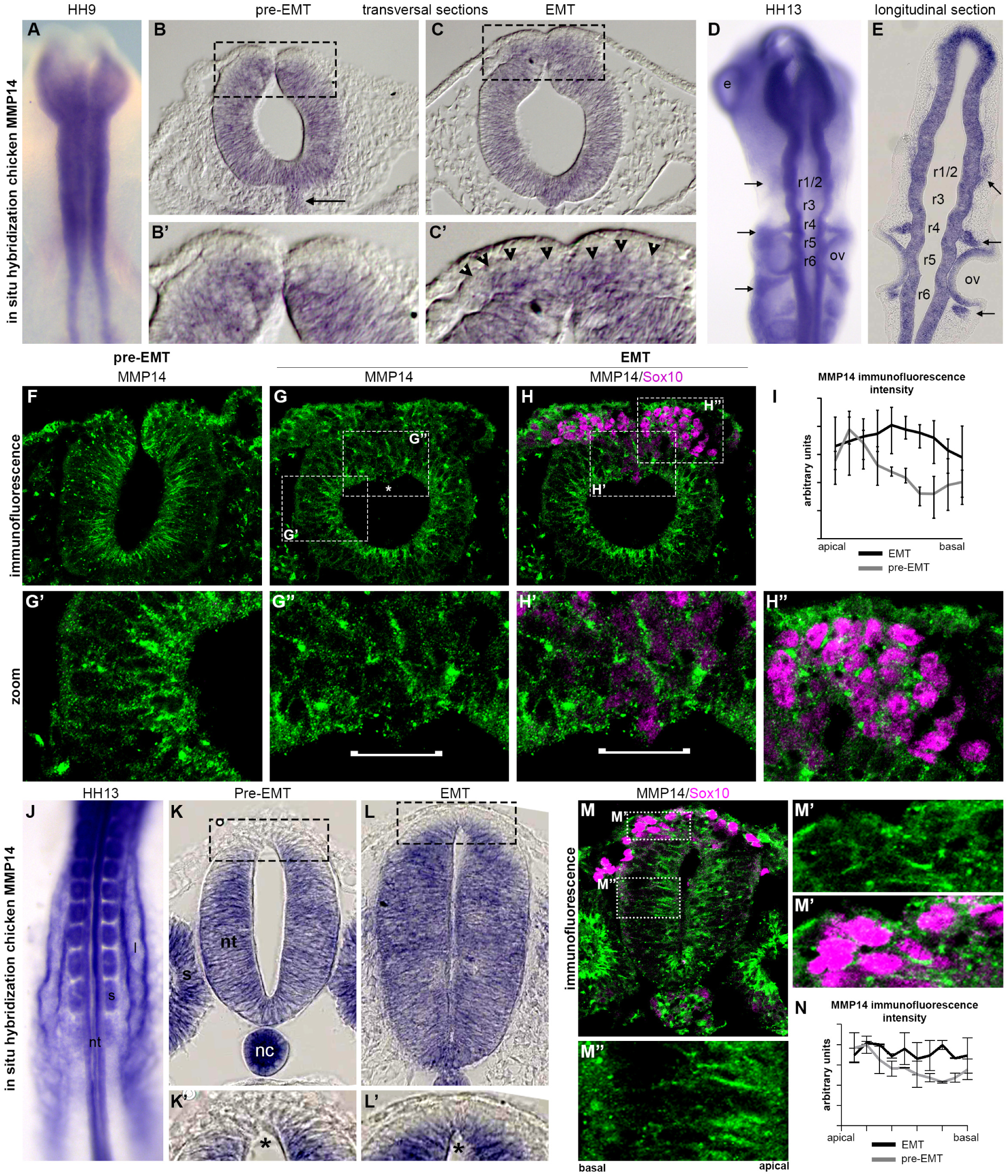
Expression and localization of MMP14 prior to and during EMT of chick NC cells.

We then monitored the subcellular localization of MMP14 protein using a custom-made mouse polyclonal antibody raised against chick MMP14 (see methods). Prior to EMT, MMP14 is found in the apical half of the neural tube (Fig. 1F, 1I grey line). At the time of NC EMT, MMP14 is localized apically in the regions outside of the NC domain (Fig. 1G-G’) while in NC cells MMP14 is more evenly distributed along the entire apicobasal axis (Fig. 1G-G’’, I, black line). On some occasion, the apical-most localization is lost (Fig. 1G, asterisk, G’, bracket) specifically in Sox10-positive NC cells (Fig. 1H, H’, bracket) suggesting that this change of subcellular distribution is directly linked to NC delamination. In addition, MMP14 is strongly detected in the cytoplasm of Sox10-positive migrating NC cells (Fig. 1H, H’’).

EMT in cephalic NC cells occurs within a short time window and nearly all cells depart at the same time concomitantly to neural folds fusion (Gouignard et al., 2018). At trunk level, the situation is different. Cells delaminate after neural tube closure and leave the neuroepithelium one by one in a dripping fashion (Gouignard et al., 2018). Thus, we wondered if the dynamics of MMP14 expression and localization might be different. By in situ hybridization, MMP14 is detected in the whole neural tube (nt), the somites (s) and the lateral mesoderm (l) (Fig. 1J). On transversal sections, MMP14 is expressed in the neural tube prior to NC EMT but expression is weak in the dorsal-most part which contains the pre-migratory NC cells (Fig. 1K, K’, asterisk). During EMT, MMP14 expression increases within the NC domain (Fig. 1L, L’, asterisk). By immunostaining, MMP14 protein is detected in the apical part of neuroepithelial cells (Fig. 1M, M’’) whereas it is broadly distributed along the apicobasal axis in NC cells (Fig. 1M, M’, N).

MMP14 is produced as an inactive precursor. It has a pro-domain in N-terminal position that acts as an endogenous catalytic inhibitor (Tallant et al., 2010). MMP14 is activated in the pathway to the plasma membrane by Furin-like pro-enzymes convertases that remove the pro-domain and is thus displayed at the cell surface in its active form. We generated a probe against chicken Furin and found it expressed in the whole neural tube (Fig. S1) suggesting that MMP14 may be processed. To substantiate this observation, we performed a western blot on total protein extracts from stage HH12 embryos and found several bands corresponding to the various forms previously described for human MMP14 (Lehti et al., 1998; Toth et al., 2002), including the full length pro-MMP14 at 68kDa, a slightly shorter band corresponding to the active form and two smaller forms which most likely correspond to the result of an auto-catalytic cleavage removing the catalytic domain (Hernandez-Barrantes et al., 2000; Lehti et al., 1998) and a secreted form obtained by shedding of the whole extracellular domain (Toth et al., 2002; Toth et al., 2005).

A-E, in situ hybridization against chick MMP14. A, Whole mount at stage HH9. B-B’, transversal cryosections from a stage HH8 embryo (mesencephalon). C-C’, transversal cryosections from a stage HH9 embryo (mesencephalon). D, whole mount at HH13. E, longitudinal cryosection from a stage HH13 embryo. F-H’’, immunostaining against chick MMP14 (green). F, cryosection from a stage HH8 embryo (mesencephalon). G-H’’, cryosections from a stage HH9+ embryo (mesencephalon) counterstained with an anti-Sox10 antibody to mark the NC cells (magenta). Asterisk and brackets indicate Sox10-positive cells that have lost apical localization of MMP14. Note that in intermediate portion of the neural tube or in the NC domain before EMT, MMP14 is mostly apical whereas in NC undergoing EMT it is found all along the apicobasal axis. I, plot of mean fluorescence intensity of MMP14 immunostaining in cephalic NC domain along the apicobasal axis at pre-EMT (grey line) and EMT stage (black line). Error bars represent standard deviation, 3 sections were used. J-L’, in situ hybridization against chick MMP14. J, dorsal view of the caudal part of a stage HH13 embryo. K-L’, cryosections. Boxes indicate the region of the zooms in K’ and L’. Note the increase of MMP14 expression at the level of pre-migratory NC cells (asterisks). M-M’’, immunostaining against chick MMP14 (green). Counterstaining against Sox10 (Magenta). M’, NC region. M’’, intermediate neural tube region. N, Plot of mean fluorescence intensity of MMP14 immunostaining in trunk NC domain along the apicobasal axis at pre-EMT (grey line) and EMT stage (black line). Error bars represent standard deviation, 3 sections were used.

Altogether, our results indicate that i) MMP14 and its main activator, Furin, are broadly expressed in whole neural tube including the NC domain, ii) that MMP14 is activated and processed and iii) that MM14 subcellular localization specifically changes in NC cells at the time of EMT.

### MMP14 is required for chick cephalic Neural Crest EMT independently of its catalytic activity

To test whether MMP14 activity is required for NC delamination, we used several independent methods for loss of function. We started with a chemical inhibitor, called NSC405020 targeting the hemopexin domain of MMP14 (Remacle et al., 2012). This inhibitor does not block the catalytic activity of MMP14 but interferes with its abilities to interact with co-factors and substrates. To apply this inhibitor, we took embryos out of their eggs at stage HH6 and folded them, a technique known as Cornish pasty culture (Nagai et al., 2011; Nagai et al., 2014). This maintains the tension required for normal development and allows to grow up to 10 embryos in 1ml of culture medium (Fig. 2A). Embryos were cultured overnight with DMSO as control or with NSC405020 (0.5mM). The DMSO did not lead to any noticeable defects. The pre-otic (rhombomere 4) and post-otic (rhombomere 6) NC streams are seen migrating normally (Fig. 2B, arrowheads) whereas in embryos treated with NSC405020 the number of migrating cells is dramatically reduced (Fig. 2C, arrowheads). Accordingly, the size of the NC domain within the neural tube (Fig. 2D) and the distance of migration (Fig. 2E) are both severely reduced. These data suggest that MMP14 is required for NC delamination. To confirm this, we designed an anti-splicing antisense Morpholino and two different siRNA (Fig S2). In addition, we made several expression vectors for various forms of MMP14 for rescue purposes: a full length MMP14 (FL), a form lacking the catalytic domain (ΔCat) and a form missing the cytoplasmic domain (ΔCyto). The catalytic activity of MMPs (or ADAMs) is heavily depending on a Glutamate located in the zinc-binding consensus sequence of the catalytic domain (Tallant et al., 2010). A simple point mutation replacing this Glutamate (E) by an Alanine (A) or a Glutamine (Q) is sufficient to abolish the catalytic activity. This has been previously showed for numerous proteases including MMP14 (Gutierrez-Fernandez et al., 2015), MMP19 (Sadowski et al., 2003), MMP28 (Maldonado et al., 2018; Rodgers et al., 2009) and ADAM13 (Khedgikar et al., 2017), Thus, we generated one form of chicken MMP14 containing a point mutation abolishing the catalytic activity by replacing E223 by an A (MMP14-EA). All these constructs are tagged in their C-terminal domain (GFP or mCherry). Importantly, these expression constructs were generated from a full length sequence of chick MMP14 obtained by codon optimization such that the siRNA vectors and the anti-splicing MO do not recognize them (see Methods).

**Figure 2.**
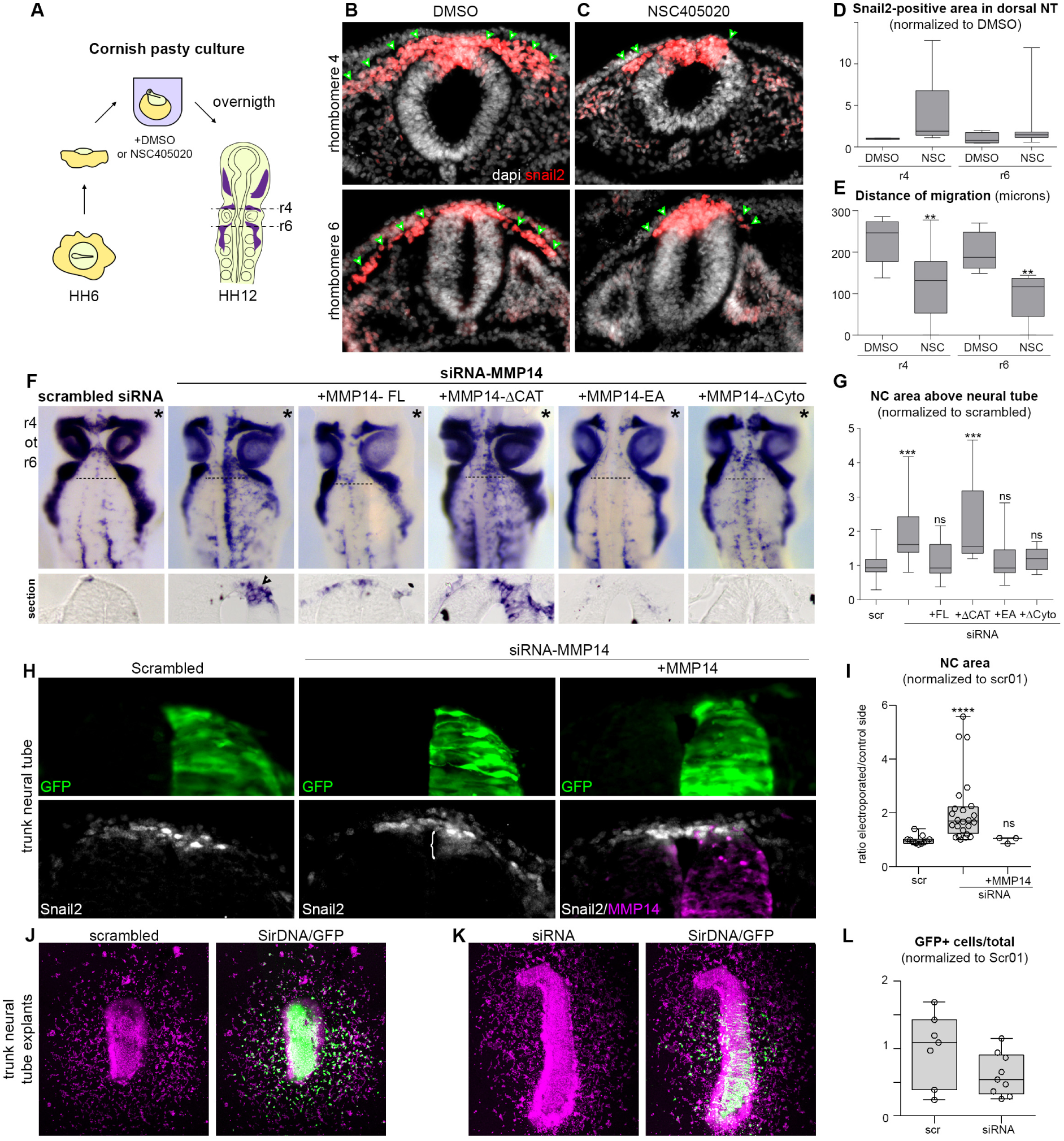
MMP14 is required chick NC EMT independently of its catalytic activity. A, diagram showing the main steps of the Cornish pasty culture technique. B-C, immunostaining for Snail2 on cryosections of embryos treated with DMSO (B) or MMP14 inhibitor NSC405020 at 0.5mM (C). D, plot showing the size of the Snail2-positive region normalized to DMSO in the dorsal neural tube (DMSO: n_embryos_=4, n_sections_=8; NSC: n_embryos_=7, n_sections_=7, from one experiment). Error bars show the standard deviation. E, distance of migration from the dorsal midline (DMSO: n_embryos_=11, n_sections_=11; NSC: n_embryos_=11, n_sections_=11, from two experiments). Error bars represent the standard deviation. ANOVA, followed by multiple comparisons; **, p value <0.01. F, in situ hybridization using a mix of probes against Foxd3 and Sox10 to stain the cephalic NC cells after electroporation of control scrambled siRNA (first column, n=28 embryos from 7 experiments), siRNA-MMP14 alone (second column, n=46 embryos from 11 experiments), siRNA-MMP14 and full length MMP14 (third column, n=17 embryos from 6 experiments), siRNA-MMP14 and the delta-catalytic form (fourth column, n=11 embryos from 3 experiments), siRNA-MMP14 and the inactive point mutant (fifth column, n=14 embryos from 2 experiments), siRNA-MMP14 and the delta-cytoplasmic form (sixth column, n=11 embryos from 3 experiments). Dotted lines indicate the level of the sections shown below each whole mount. G, plot of the area occupied by Foxd3/Sox10-positive NC cells above the post-otic neural tube from whole mount images. ANOVA with Kruskal-Wallis multiple comparisons, ***p<0.0005. H, immunostaining against Snail2 after electroporation of scrambled siRNA, siRNA-MMP14 and co-electroporation with siRNA-MMP14 and full length MMP14. Scrambled and siRNA are in green (GFP), MMP14-FL-mCherry is in magenta. I, plot of the NC area (Snail2) with scrambled (n_embryos_=11, n_sections_=132), siRNA-MMP14 (n_embryos_=24, n_sections_=303) and siRNA-MMP14 and full length MMP14 (n_embryos_=3, n_sections_=15), ANOVA with Kruskal-Wallis multiple comparisons, ****p<0.0001. J-K, neural tube explants cultured on fibronectin coated dishes counterstained with SiR-DNA. Cells electroporated with scrambled siRNA or SiRNA-MMP14 are GFP-positive. L, ratio of GFP-positive cells over the total number of cells counted from SiR-DNA staining, n_scrambled_=7 explants, n_siRNA MMP14_=9 explants from 4 independent experiments. Unpaired t-test with Welsh’s correction, p=0.1186.

We electroporated these different constructs in the cephalic region and analyzed their effects 18 hours later in the post-otic domain (Fig. 2F-G, asterisks mark the electroporated side). SiRNAs and MO efficiencies on MMP14 expression were first validated (Fig. S2). The two siRNA and the splicing MO inhibited NC delamination as seen by the accumulation of Foxd3/Sox10-positive cells along the dorsal midline (Fig. 2F second column, Fig. S2). Electroporation of a scrambled siRNA sequence did not affect NC delamination (Fig. 2F, first column) whereas siRNA-MMP14 led to an accumulation of NC cells in the dorsal neural tube on the electroporated side (Fig. 2F, second column; section, arrowhead).

To decipher whether MMP14 acts via its catalytic activity, we attempted to rescue the effect of the siRNA by co-electroporating the various forms of MMP14. Strikingly, the full-length form (Fig. 2F, +MMP14-FL, third column), the inactive point mutant (Fig. 2F, +MMP14-EA, fifth column) and the delta-cytoplasmic version (Fig. 2F, +MMP14-ΔCyto, sixth column) were equally efficient at rescuing MMP14 loss-of-function (Fig. 2F-G). However, the delta-catalytic form was unable to rescue the siRNA (Fig. 2F, fourth column, section, arrowheads, 2G). These data indicate that the extracellular domain of MMP14 is required for its function during NC delamination but that its catalytic activity is not.

We next assessed whether MMP14 was also required in the trunk region where EMT is more progressive. We electroporated the scrambled siRNA in the right-hand side of stage HH12 embryos but found no effect on Snail2-positive cells (Fig. 2H, first column, 2I). However, electroporation of the siRNA-MMP14 increased the Snail2-positive domain in the neural tube (Fig. 2H, second column, bracket; 2I), an effect that is rescued by co-electroporating the full length MMP14 (Fig. 2H, third column, 2I). We then cultured neural tube explants after electroporation on fibronectin (Fig. 2J-K) and found that the siRNA-MMP14 specifically reduced the proportion of GFP-positive cells that had delaminated (Fig. 2K-L) compared to the scrambled siRNA (Fig. 2J, 2L). This indicate that blocking MMP14 is the trunk neural tube prevents EMT of NC cells and that this effect is likely to be cell autonomous since GFP-negative cells delaminated and migrated away from the neural tube. Altogether, these experiments demonstrate that MMP14 is equally required during the intense cephalic NC EMT and the slow-paced trunk NC EMT.

### Ets1 is sufficient to induce expression and basolateral localization of MMP14

The timing of chick cephalic NC delamination is controlled by the loss of p53 expression that triggers an increase of Snail2 and Ets1 expressions (Rinon et al., 2011). Interestingly, p53 is known as a transcriptional repressor of MMP14 (Cathcart et al., 2016) and Ets1 and Snail2 were shown to induce MMP14 expression in other cell types (Heo and Cho, 2014; Shields et al., 2012; Welch-Reardon et al., 2014). We thus checked whether Ets1 or Snail2 were able to upregulate MMP14 in the chick neural tube. We found that Ets1 overexpression was sufficient to induce MMP14 (Fig. 3A, arrowhead) whereas Snail2 was not (Fig. 3B). As previously mentioned, there are examples of MMP14-dependent regulation of gene expression (Shimizu-Hirota et al., 2012). We wanted to test if MMP14 might be involved in a feedback loop controlling Ets1 or Snail2 expression. Loss of MMP14 affected neither Ets1 nor Snail2 expressions in the dorsal neural tube 18h after electroporation (Fig. S3).

**Figure 3.**
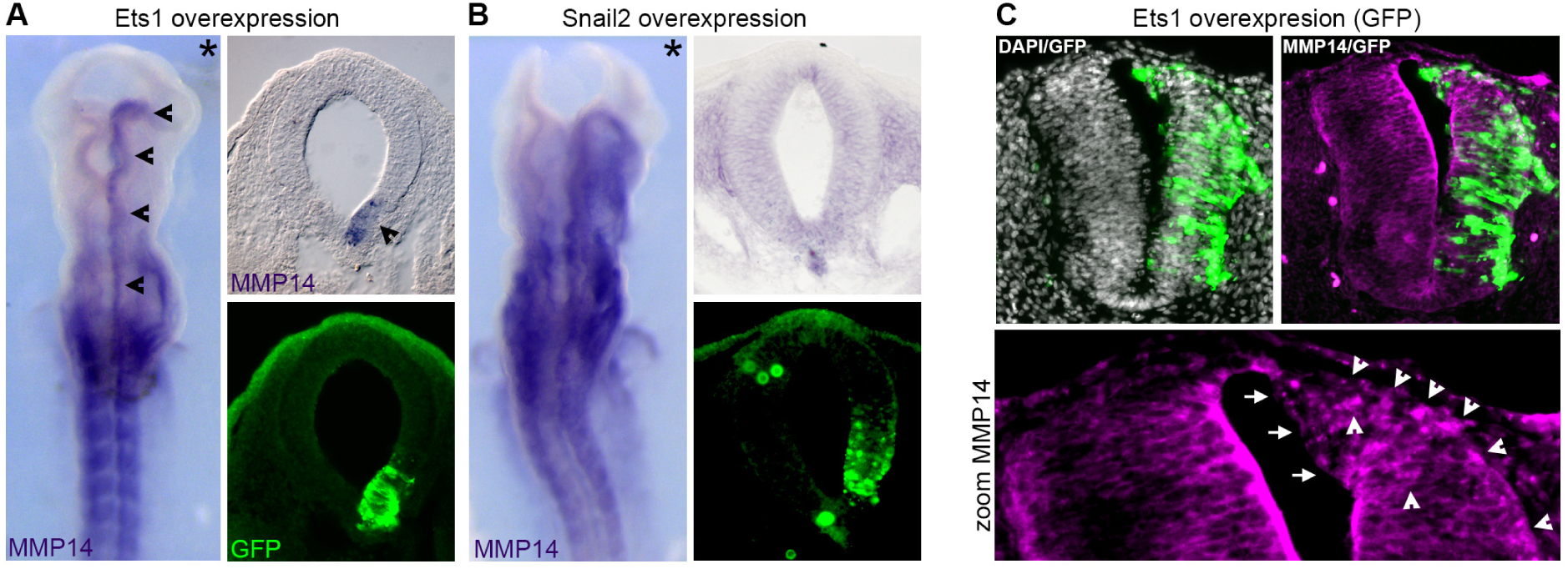
Ets1 induces expression and basolateral localization of MMP14. A-B, in situ hybridization for chick MMP14 after Ets1 overexpression (A) or Snail2 overexpression (B) in cephalic regions, n=5 embryos each. Arrowhead indicates ectopic activation of MMP14. C, Immunostaining for MMP14 (magenta) after Ets1 overexpression (GFP) in the trunk. Nuclei are stained with DAPI. Arrows indicate the loss of apical MMP14, arrowheads indicate regions of basolateral accumulation of MMP14, n=3 embryos.

MMP14 expression precedes that of Ets1 in the NC domain and MMP14 is expressed broadly in the neural tube whereas Ets1 is not. Thus, one cannot simply propose that Ets1 would trigger EMT of cephalic NC cells by inducing MMP14 expression. We wondered instead whether expressing Ets1 might be sufficient to drive a change of subcellular localization of MMP14. To test that we expressed Ets1 in the dorsal region of trunk neural tubes and performed an immunostaining for MMP14 (Fig. 3C). Interestingly, Ets1 expression was sufficient to induce a loss of apical localization of MMP14 (Fig. 3C, arrows) and an accumulation of MMP14 in basolateral positions (Fig. 3C, arrowheads). This results suggest that the peak of Ets1 expression at the onset of cephalic NC EMT might contribute to the change of apicobasal distribution of MMP14.

### MMP14 is required for the loss of cadherin-6B in pre-migratory NC cells

At the time of delamination, NC change cadherins and activate a migratory program. Thus, to understand the molecular mechanism of the delamination defect induced by the loss of MMP14, we investigated the effect of MMP14 knockdown on genes downstream of the transcription factors controlling EMT in cephalic NC cells. We analyzed Cadherin-6B (Fig. 4A), Cadherin-7 (Fig. 4B), the small GTPase RhoB (Fig. 4C) and Integrin-β3 (Fig. 4D) by in situ hybridization. We found no effect on Cadherin-7 (Fig. 4B) whereas the area of Cadherin-6B expression was extended after inhibition of MMP14 (Fig. 4A). In addition, cells that are retained in the dorsal neural tube express RhoB (Fig. 4C) and Integrin-β3 (Fig. 4D). Detecting RhoB-positive/integrin-β3-positive cells in the dorsal neural tube after MMP14 knockdown strongly suggest that cells are adopting a migratory phenotype. However, the maintenance of Cadherin-6B and the absence of Cadherin-7 in the dorsal neural tube indicate that the switch from 6B to 7 did not occur in cells electroporated by siRNA-MMP14.

**Figure 4.**
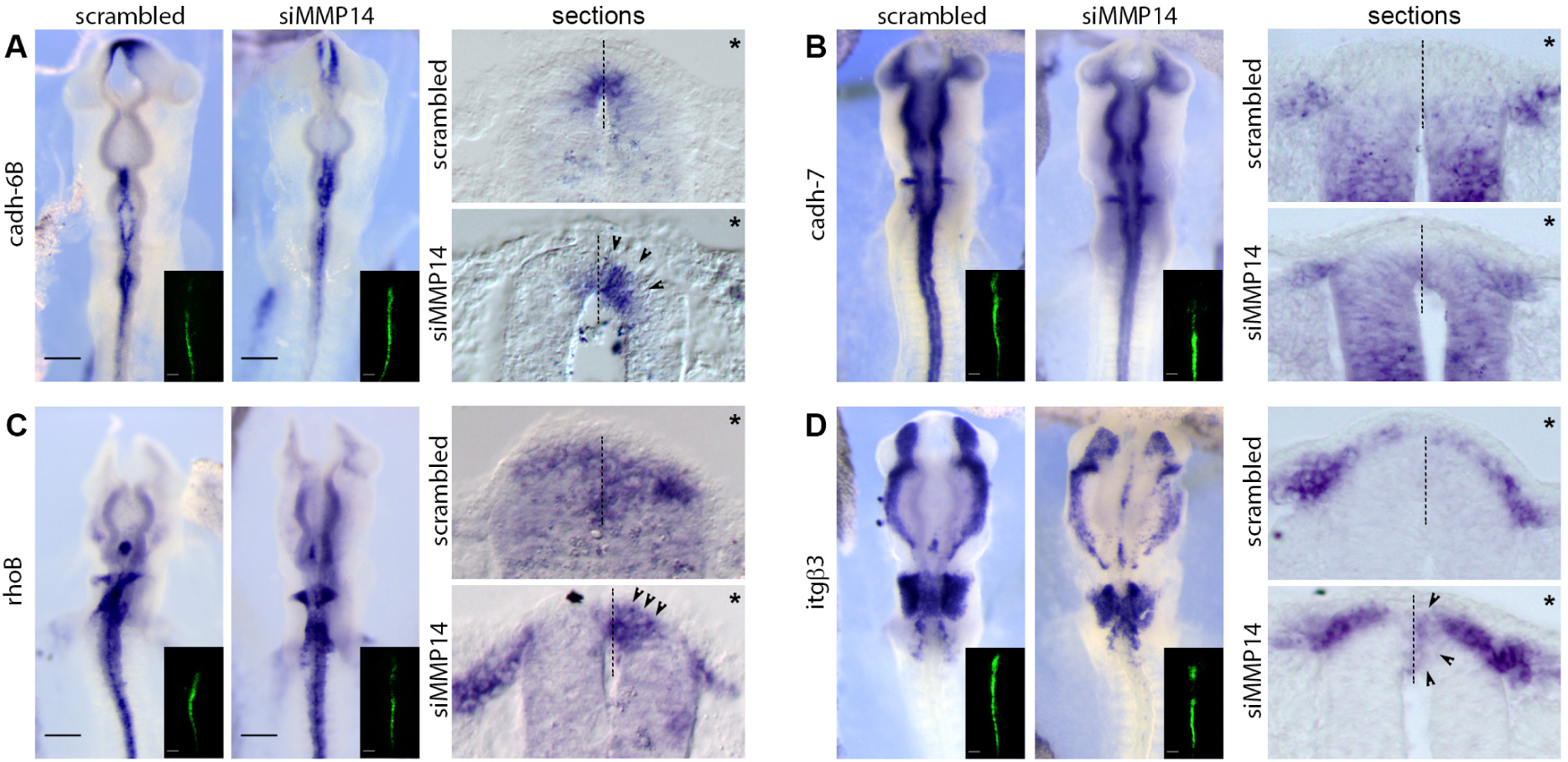
MMP14 is required for the loss of Cadherin-6B. A-D, in situ hybridization and transversal cryosections through the post-otic NC region for Cadherin-6B (A), Cadherin-7 (B), RhoB (C) and Integrin-β3 (D) in embryos electroporated with scrambled (left whole mount, top section) or MMP14 (right whole mount, bottom section) siRNA, n_scrambled_=23, n_siRNA-MMP14_=22 ; from 2 experiments. Asterisks on sections mark the electroporated side, insets in whole mount images indicate the electroporated regions. Dotted line delineate the midline of the neural tube. Arrowheads indicate NC cells located within the dorsal neural tube expressing the gene of interest.

### MMP14 is required for the loss of apicobasal polarity in pre-migratory NC cells

We then looked at E and N-cadherin expressions by immunostaining. Knocking down MMP14 had no effect on their distributions (Fig. 5A-D). Next, we stained for Pericentriolar Material 1 (PCM1, Fig. 5E-F’). PCM1 is apically located in the neural tube prior to EMT and then redistributed throughout the cells during EMT (Fig. S4). Thus, it is a good marker to assess changes in apicobasal polarity in NC cells. The scrambled siRNA did not affect the typical change of subcellular localization of PCM1 (Fig. 5E) as all cells located in the dorsal-most neural tube domain have PCM1 in non-apical locations (Fig. 5E, arrowheads) on the control and the electroporated sides. However, siRNA-MMP14 prevented the delocalization of PCM1 along the basolateral membrane (Fig. 5F). Indeed, AP2-positive cells that are retained within the electroporated half of the neural tube have a clear apical localization of PCM1 (Fig. 5F, bracket; F’ right-hand side of the image) by contrast to the diffuse distribution observed on the contralateral non-electroporated side (Fig. 5F-F’, arrowheads). This result indicates that MMP14 is required for the change of polarity during NC EMT.

**Figure 5.**
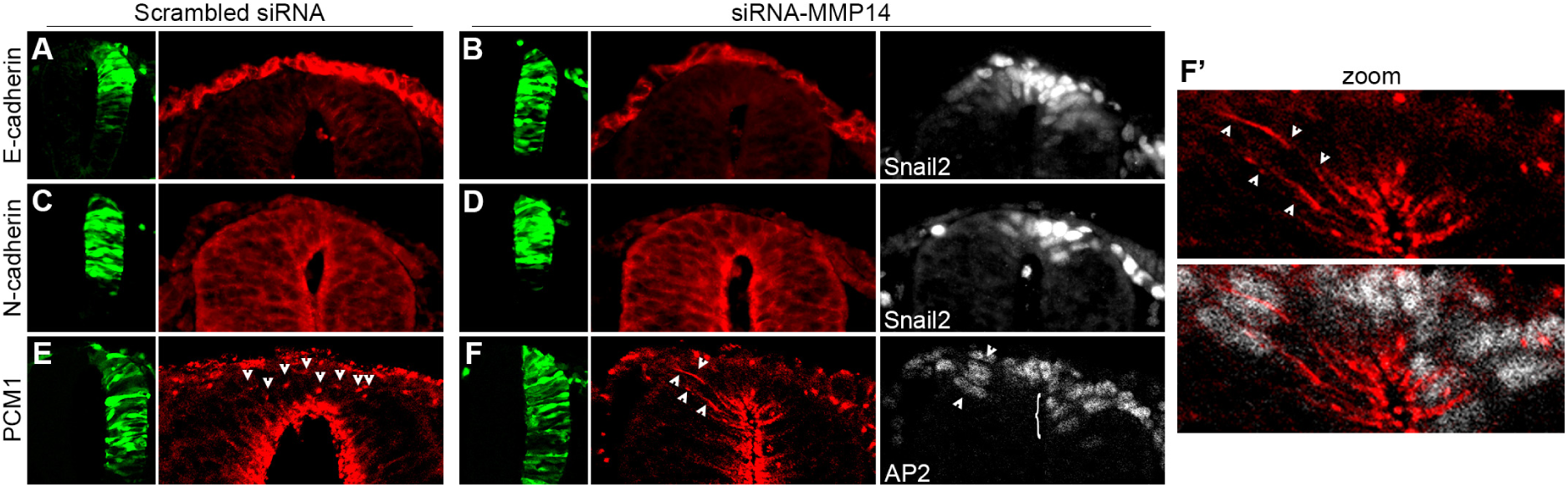
MMP14 is required for the loss of apicobasal polarity. A-B, immunostaining against E-cadherin (red) and Snail2 (white) in embryos electroporated with scrambled (A, n=5 embryos) or MMP14 (B, n=3 embryos) siRNA. C-D, immunostaining against N-cadherin (red) and Snail2 (white) in embryos electroporated with scrambled (C, n=5 embryos) or MMP14 (D, n=3 embryos) siRNA. E-F’, immunostaining against pericentriolar material 1 (PCM1) and AP2 (white) in embryos electroporated with scrambled (E, n=6 embryos) or MMP14 (F-F’, n=5 embryos) siRNA. Arrowheads in E indicate basolateral localization of PCM1 on both the non-electroporated and the scrambled-siRNA sides. Arrowheads in F marks the basolateral localization of PCM1 on the non-electroporated side only. Note that PCM1 is solely found within the apical domain in AP2-positive cells after electroporation of siRNA-MMP14 (bracket). F’, zoom on the neural crest domain. Electroporated cells are visualized by GFP expression (green). Data from 2 independent experiments.

### Basolateral localization of MMP14 is sufficient to affect apicobasal polarity

We then wondered whether MMP14 needed to be removed from its apical localization or to be sent to basolateral positions to affect apicobasal polarity. In the cephalic regions, overall polarity is difficult to study at the time of EMT because all cells are undergoing EMT simultaneously. In the trunk, neuroepithelial cells undergo mitosis in the apical domain. This is linked to a strong apicobasal polarity and can be affected directly by modulating the localization of apical markers such as Par3 (Afonso and Henrique, 2006). In the NC domain, this strict apical localization of mitoses is lost due to the loss of apicobasal polarity during EMT and a small fraction of NC cells perform basal mitosis prior to exiting the neural tube (Ahlstrom and Erickson, 2009). Thus, one can use position of mitotic nuclei to assess the polarity of cells in the neuroepithelium and in the neural crest domain. We first electroporated the scrambled and the MMP14 siRNA and found that only siRNA-MMP14 reduced the proportion of basally dividing NC cells (Fig. 6A-B). This result indicate that in absence of MMP14 NC cells behave like control neuroepithelial cells and position their nuclei apically for mitosis. Next, we analyzed the effect of knocking down MMP14 in neuroepithelial cells. Endogenous MMP14 is apical in the non-NC regions of the neuroepithelium (Fig. 1M, M’’). Neither the scrambled siRNA nor the MMP14 siRNA had any effect on the overall localization of mitotic nuclei (Fig. 6C-D). These data indicate that apical localization of MMP14 is not required for normal apicobasal polarity of neuroepithelial cells.

**Figure 6.**
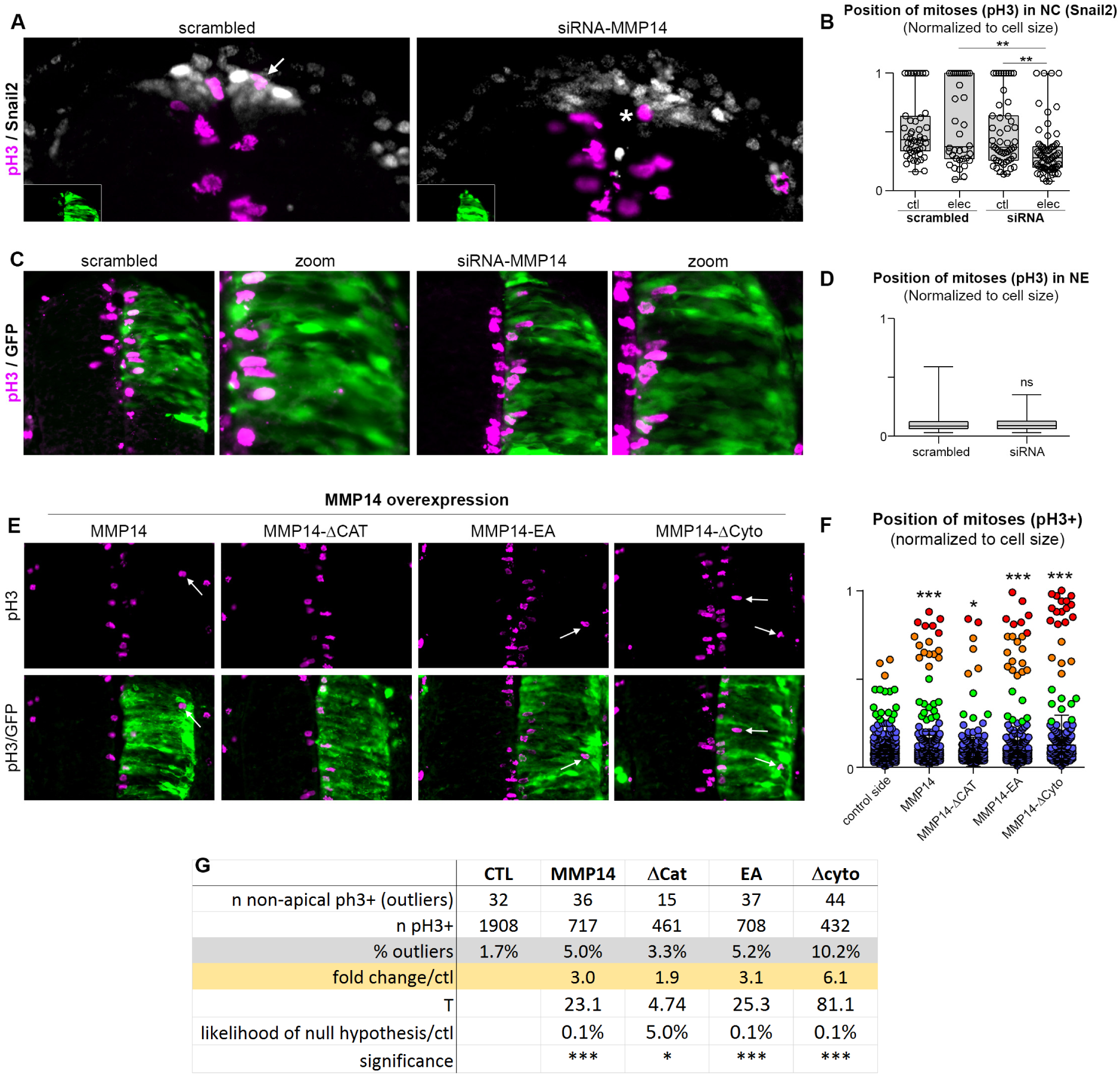
Basolateral localization of MMP14 is sufficient to impair apicobasal polarity independently of its catalytic activity.

A, immunostaining against phosphor-histone H3 (pH3, magenta) and Snail2 (white), zones of electroporation are shown in insets on the bottom left corner of each image. Arrow shows an example of basal mitosis in NC cells electroporated with scrambled siRNA. Asterisk marks an example of an apically located mitosis in NC cells electroporated by the siRNA-MMP14. B, position of mitoses in NC cells on the control non-electroporated side and scrambled or MMP14 siRNA electroporated side for each condition. Note that in non-electroporated cells and scrambled cells there is a sub-population of basal mitoses (clustering at 1). This sub-population is dramatically reduced in siRNA-MMP14 cells. ANOVA followed by Kruskal-Wallis multiple comparisons; **(scr/si), p=0.0011 ; **(non-electroporated si/si), p=0.0024. Scrambled siRNA, n=3 embryos (36 mitoses), siRNA-MMP14, n=8 embryos (69 mitoses), non-electroporated cells, n=11 embryos (99 mitoses) from 3 experiments. C-D, immunostaining for phosphor-histone H3 (pH3, magenta) after electroporation of scrambled (left panels) and MMP14 (right panels) siRNA. GFP corresponds to electroporated cells. D, position of mitoses in neuroepithelial cells (intermediate regions of the neural tube) on the electroporated side of scrambled (n=3 embryos (197 mitoses)) or MMP14 (n=9 embryos (270 mitoses)) siRNA embryos, from 3 experiments. Mann-Whitney test, p=0.8391. E, immunostaining for phospho histone H3 (pH3, magenta) after overexpression of, from left to right, MMP14, MMP14-ΔCat, MMP14-EA, MMP14-ΔCyto. GFP corresponds to electroporated cells. Arrows indicate mitoses occurring away from the apical side. F, Position of mitoses for each condition depicted in E. All mitoses from the non-electroporated sides were pooled. Note that there are two populations of mitotic nuclei, one apical (shades of cold colors, clustering near 0) and one away from the apical domain (shades of warm colors). G, Proportions of non-apical mitoses (outliers) for each conditions shown in E and F and within the non-electroporated epithelium. Statistics for proportions were performed according to (Taillard et al., 2008). Thresholds for T were such that at T > 3.841 the null hypothesis is rejected with a 95% confidence (*), at T > 6.635 the null hypothesis is rejected with a 99% confidence (**), at T > 10.83 the null hypothesis is rejected with a 99.9% confidence (***). Data from 4 independent experiments.

To test whether basolateral localization of MMP14 would be sufficient to affect polarity, we overexpressed various forms of MMP14 in the neuroepithelium (Fig. 6E). All forms of MMP14 (FL, EA, ΔCat, Δcyto) were able to produce basal mitoses (Fig. 6E-F, arrows). We defined ectopic mitoses (outliers) as cell division occurring more than two nuclei diameter away from the apical domain. With that criteria, we observed 1.7% of ectopic mitoses in non-electroporated control sides from all conditions. Expressing full length MMP14 or the inactive point mutant raised this category to 5% of the total number of mitoses. Interestingly, the ΔCat version was less efficient and produced 3.3% of ectopic mitoses whereas the ΔCyto form was the most efficient with 10% of basal mitoses (Fig. 6G). Proportions of non-apical mitoses in all conditions were statistically compared to controls (Taillard et al., 2008) (Fig. 6G).

A, Diagram summarizing the various gain and loss of function experiments performed in NC cells and neuroepithelial cells and their impact on the position of mitotic nuclei. Nuclei are depicted in grey but are not represented as performing mitoses. MMP14 is in green, N-cadherin in red, PCM1 in blue, laminin in brown. +++, normal apicobasal polarity; +/- impaired apicobasal polarity. B, Simplest model for the role of MMP14 in EMT from the data presented in the present study. MMP14 is placed downstream of pro-EMT genes and upstream of apicobasal polarity. Red lines indicate negative regulation. KD, knockdown; OE, overexpression.

Next, we wanted to check whether forced basolateral expression of MMP14 would be sufficient to lead to EMT-like phenotype such as loss of N-cadherin, degradation of Laminin or apical detachment of neuroepithelial cells. We found no evidence of such defects (Fig. S5) indicating that the basal mitoses induced by MMP14 are not a consequence of cells undergoing delamination in ectopic locations of the neural tube. Instead, they are most likely due to a direct effect of MMP14 on apicobasal polarity.

## Discussion

In this study, we used the NC cells and the neuroepithelium of the chick embryo to study the role of MMP14 is physiological EMT. We found that while its expression is not restricted to cells undergoing EMT or migration, the protein displays a dynamic change of subcellular localization during EMT. We found that it primarily acts via a destabilization of apicobasal polarity while located in basolateral domains of the cells. In neuroepithelial cells, or pre-EMT NC cells, MMP14 is apical. Loss of function does not affect polarity in neuroepithelial cells but maintains normal apicobasal polarity in NC cells performing EMT. These data strongly suggests that, while it is in apical localization, MMP14 is stored and plays no direct role in apicobasal polarity. By contrast its basolateral localization during EMT is essential for the change of cell polarity. These has been demonstrated by gain and loss of functions and is summarized in Fig. 7A. Overall, MMP14 seems to act downstream of EMT regulator genes (e.g. Ets1) without any obvious feedback loop from MMP14 to these genes (Fig. 7B).

**Figure 7.**
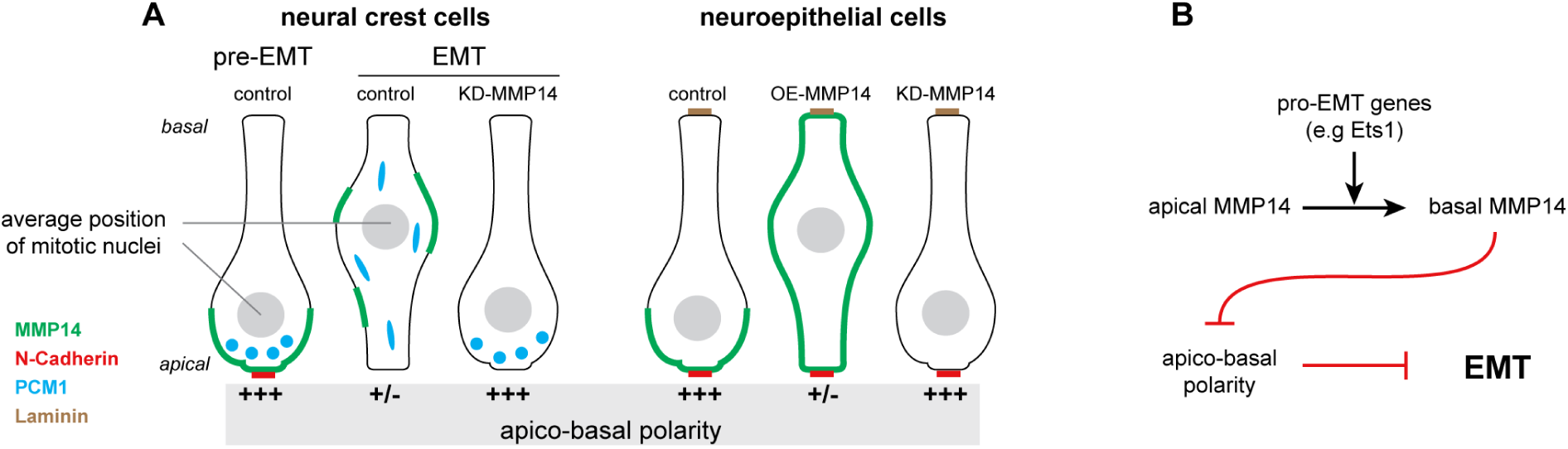
Basolateral localization of MMP14 contributes to EMT by destabilizing apicobasal polarity.

Literature on cephalic NC EMT has demonstrated a key role of Cadherin-6B in cephalic NC EMT (Coles et al., 2007; Schiffmacher et al., 2014; Schiffmacher et al., 2016; Taneyhill et al., 2007). Surprisingly, we showed that MMP14 is required for the loss of Cadherin-6B expression. While MMP14 was previously shown to traffic to the nucleus and display some transcriptional activity (Shimizu-Hirota et al., 2012), we never observed MMP14 in the nucleus by immunostaining. In addition, Cadherin-6B expression is known to be downregulated by Snail2 (Taneyhill et al., 2007). In our MMP14 knockdown the domain of Snail2-positive cells within the dorsal neural tube is enlarged indicating that the loss of Cadherin-6B repression is not due to a loss of Snail2. These observations, in the context of the current literature on cephalic NC EMT, strongly suggest that MMP14/Cadherin-6B interaction at transcriptional and protein levels is unlikely to be direct and will need to be further studied. While a direct of Cadherin-6B in apicobasal polarity has been demonstrated, it is known that preventing its cleavage by ADAMs maintains its apicolateral localization in pre-migratory NC cells (Schiffmacher et al., 2014). Thus an enhanced level of Cadherin-6B might contribute to a stronger apicobasal polarity simply by tilting the Cadherin/ADAM balance in favor of Cadherin-6B. However, it is important to note that the role of MMP14 in apicobasal polarity is not bound to Cadherin-6B expression. Indeed, basolateral expression of MMP14 in intermediate neuroepithelial cells was sufficient to impair polarity and generate basal mitoses. These cells do not express Cadherin-6B and therefore MMP14 is able to modulate polarity independently of Cadherin-6B. In addition, whereas cephalic NC cells delaminate as Cad-6B-/MMP14+, trunk NC cells delaminate as Cad-6B+/MMP14+ indicating that there is no systematic inverse correlation between the expressions of the two genes.

MMP14 expression and function are often discussed in the context of invasion due to its ability to activate another collagenase called MMP2 (Sato et al., 1994). However, our results show that MMP14 is not restricted to sites of EMT and migration. This is in sharp contrast with previously published expression patterns for MMP2 in the early chicken embryo clearly linking MMP2 expression with EMT (Duong and Erickson, 2004). However, functions of MMP14 in tissues that do not express MMP2 (e.g. early epithelial somites, neural tube) may still partially depend on MMP2. MMP2 is a secreted protein and cells expressing MMP14 at their cell surface may activate MMP2 produced by another cell population. Such model has been proposed for the migration of cephalic NC cells where migratory cells expressing MMP14 might activate latent MMP2 found in the mesenchyme (Cantemir et al., 2004). In Xenopus the requirement of MMP14 in NC cells and MMP2 in the local environment for successful NC migration seems clear, however activation of MMP2 in this context may not require MMP14 (Garmon et al., 2018).

MMP2 is usually activated by a dimer of MMP14 associated with one molecule of tissue inhibitor of metalloproteinase 2 (TIMP2) (Goldberg et al., 1989; Murphy et al., 1992). Interestingly, chicken MMP14 lacks a domain called the MT-loop (Yang et al., 2008) that is critical for the activation of MMP2 (English et al., 2001). In addition, our results indicate that MMP14 acts independently of its catalytic activity strongly suggesting that MMP14 and MMP2 may not belong to the same pathway in the chick embryo. Other MMPs would have to activate MMP2 instead such as MMP15/MT2-MMP (Morrison et al., 2001), MMP16/MT3 (Nakada et al., 1999) which are expressed by cephalic NC cells at the time of EMT (Schiffmacher et al., 2018). Intriguingly, Cadherin-6B can modulate MMP2 activity (Schiffmacher et al., 2018). Cadherin-6B is processed by ADAM proteases and a soluble N-terminal fragment (NTF) is released. It was shown that this fragment is sufficient to promote an increase of MMP2 activity. In vitro, a physical interaction has been found between MMP2 and Cadherin-6B potentially stabilizing MMP2. Given that MMP2 can self-activate by autocatalysis of its pro-domain (Lafleur et al., 2003) such stabilization of MMP2 by physical interaction with Cadherin-6B may promote MMP2 activation independently of other pro-protein convertases.

In endothelial cells in culture, MMP14 is seen at cell-cell contacts associated with integrin-β1 whereas it goes to protrusions and associates with integrin-β3 upon wounding (Galvez et al., 2002). In these cells, association with integrin-β1 is linked to a low turnover and a low proteolytic activity whereas co-localization with integrin-β3 is associated with a high turnover and a high proteolytic activity. Cephalic NC cells turn on integrin-β3 expression when they start their migration (Pietri et al., 2003). Therefore, one possibility would be that the change of subcellular localization from apical to basal that we observed is linked to the onset of integrin-β3 expression. In our MMP14 loss of function experiment, NC cells that fail to delaminate nonetheless express integrin-β3 while still located within the dorsal neural tube. It would be interesting to monitor the subcellular localization of integrin-β3 in control and MMP14 knockdown NC cells to see if integrin-β3 can be properly localized in the basal domain in absence of MMP14.

Human MMP14 has a cytoplasmic tail that contains several binding sites for regulators of the small GTPases Rac1 and RhoA (Gonzalo et al., 2010a; Hoshino et al., 2009) that could explain its role in cell polarity. However, some of these sites are absent in the chicken protein (Yang et al., 2008). Moreover, we showed that the form of MMP14 lacking its cytoplasmic domain was capable of rescuing the loss of function and that it was sufficient to promote basal mitoses indicating that this domain is not critical for neural crest delamination and the modulation of apicobasal polarity in chick (Fig. 2I-J). Altogether, these observations make it unlikely that chicken MMP14 would modulate polarity via direct regulation of small GTPases via its cytoplasmic domain. Instead, given that the extracellular domain is essential, one should look for extracellular or cell surface proteins interacting with MMP14 that might be involved in maintenance of apicobasal polarity.

Given that expression of MMP14 is expressed in numerous epithelial tissues in the embryo (e.g. neuroepithelium, placodes, endoderm) and in the adult (e.g. gut, lung, gonads), our results about non-catalytic functions in apicobasal polarity in vivo strongly invite to a thorough assessment of the functions of MMP14 in these tissues during homeostasis and diseases.

## Acknowledgements

The authors would like to thank Dr. Andreas Merdes (CNRS UMR5547, Toulouse) for the kind gift of the anti-PCM1 antibody (Dammermann and Merdes, 2002). We are grateful to Drs. Hervé Acloque for critical reading of the manuscript and Céline Cougoule for friendly advice throughout the project.

## Author contributions

E.T designed the project; C.A. performed most of the experiments; A.M and C.A performed the molecular biology; D.A and C.A raised the anti-MMP14 antibody; C.A. and E.T prepared the figures and supplementary information; E.T wrote the article with inputs from all authors.

## Materials and Methods

### Chicken Eggs

Fertilized chicken eggs were incubated at 38°C until the desired stage (Hamburger and Hamilton, 1992).

### Electroporation

Embryos at HH8 or HH12 were windowed. Using a glass capillary, a solution containing 6% sucrose, 0.05% Fast Green and the expression vectors of interest was injected into the lumen of the anterior neural tube and deposited onto the open neural plate. A series of seven electric pulses (28V, 5Hz) is applied to electroporate the neural tube. A few drops of PBS1X containing an antibiotic are added to the egg before being closed and placed in the incubator overnight.

### Cornish pasty culture

Embryos at stage HH6 were taken out of their egg with their extracellular tissues and placed in a dish containing Albumen/Pannett-Compton (2:1). The embryos were folded in two along the anteroposterior axis of the embryo, keeping the embryo at the surface. With a scalpel, the extremities of the extraembryonic tissues were cut off. This procedure immediately seals off the tissues together generating a small flat crescent-shaped structure with the embryo at the edge resembling a Cornish pasty (Nagai et al., 2011; Nagai et al., 2014). Pasties can then be kept floating in culture medium overnight.

### Antibody against chicken MMP14

Full length Chick MMP14-Flag was expressed in human Hek293T cells using Extreme gene HP (Sigma) and purified with anti-Flag-agarose beads (Sigma). Balbc mice (Jackson lab) were immunized with 100 µl of anti Flag-agarose beads bound to cMMP14 using complete and the incomplete freund adjuvant. Tail bleeds were collected after 2 boosts and were tested against Hek293T cells transfected with either RFP-Flag (negative control) or cMMP14.

### Histology

Embryos were soaked in Phosphate Buffer 15% sucrose overnight at 4°C. Then, embryos were transferred for 2 hours in gelatin 7.5%/ sucrose 15%. Small weighing boats are used as molds. A small layer of gelatin/sucrose is deposited at the bottom and left to set. Embryos are then transferred on the gelatin layer using a plastic 2.5mL pipette. Each embryo is placed in a single drop and left to set. Once all drops are set, an excess of gelatin/sucrose solution is poured on to the weighing boat to fill it. Once again gelatin is left to set on the bench. After setting, the dish is placed at 4°C for 1 hour to harden the gelatin. Once ready, the block of gelatin containing the embryos is placed under a dissecting microscope and individual blocks are carved to position the embryos in the desired orientation for sectioning.

### Morpholinos, siRNA and MMP14 inhibitor

Control MO 5’ CCTCTTACCTCAGTTACAATTTATA 3’, antisplicing Morpholino MMP14 5’-CCCCAGCCCCACACCTTGAACA-3’, Mo5Mismatch: 5’ CC<u>G</u>CA<u>C</u>CCC<u>G</u>ACACCTT<u>C</u>AA<u>G</u>A 3’. All siRNA and their scrambled controls were cloned into pGFR RNAi vector. The target sequence, the primers used and the final siRNA sequence are given below for each construct.

siRNA-MMP14-01 targets the following sequence 5’-GGAAGTGTCGACCCGGAAA -3’, For 1: 5’-gagaggtgctgctgagcgAGAAGTGTCGACCCGGAAATAGTGAAGCCACAGATGTA-3’, Rev 1: 5’-attcaccaccactaggcaGGAAGTGTCGACCCGGAAATACATCTGTGGCTTCACT-3’; Final sequence siRNA-MMP14-01: 5’-ggcggggctagctggagaagatgccttccggagaggtgctgctgagcg AGAAGTGTCGACCCGGAAATAGTGAAGCCACAGATGTATTTCCGGGTCGACACTTCCtgcctagtggtg gtgaatagcggggttagaagctttcttcccctcttcttaagccaccc-3’

siRNA-MMP14-02 targets the following sequence 5’-AAGGACGGCAAGTTCGTCTTC-3’, For 2: 5’-gagaggtgctgctgagcgCAGGACGGCAAGTTCGTCTTCTAGTGAAGCCACAGATGTA-3’ Rev 2: 5’-attcaccaccactaggcaAAGGACGGCAAGTTCGTCTTCTACATCTGTGGCTTCACT-3’ Final sequence siRNA-MMP14-02: 5’ggcggggctagctggagaagatgccttccggagaggtgctgctgagcgCAG GACGGCAAGTTCGTCTTCTAGTGAAGCCACAGATGTAGAAGACGAACTTGCCGTCCTTtgcctagtggtg

gtgaatagcggggttagaagctttcttcccctcttcttaagccaccc-3’

MMP14 scrambled 1 targets the following sequence: 5’-ACGGACTAGCTAAGGACGG-3’, scrFor 1: 5’-gagaggtgctgctgagcgCCGGACTAGCTAAGGACGGTAGTGAAGCCACAGATGTA-3’ scrRev 1: 5’-attcaccaccactaggcaACGGACTAGCTAAGGACGGTACATCTGTGGCTTCACT-3’ Final sequence scrambled-01: 5’-ggcggggctagctggagaagatgccttccggagaggtgctgctgagcgCCG GACTAGCTAAGGACGGTAGTGAAGCCACAGATGTACCGTCCTTAGCTAGTCCGTtgcctagtggtggtg aatagcggggttagaagctttcttcccctcttcttaagccaccc-3’

MMP14 scrambled 2 targets the following sequence: 5’-GCATCGTAATGCGCGAGTCAT-3’ scrFor 2: 5’-gagaggtgctgctgagcgACATCGTAATGCGCGAGTCATTAGTGAAGCCACAGATGTA-3’ scrRev 2: 5’-attcaccaccactaggcaGCATCGTAATGCGCGAGTCATTACATCTGTGGCTTCACT-3’ Final sequence scrambled-02: 5’-ggcggggctagctggagaagatgccttccggagaggtgctgctgagcgACA TCGTAATGCGCGAGTCATTAGTGAAGCCACAGATGTAATGACTCGCGCATTACGATGCtgcctagtg gtggtgaatagcggggttagaagctttcttcccctcttcttaagccaccc-3’. MMP14 inhibitor NSC 405020 (EMD Millipore, 444295).

### Histology and immunostainings

Gelatin embedding and cryosections using a cryostat Leica CM1950 were performed as previously described (Theveneau 2007). Sections were incubated in PBS1X at 42°C for 30 minutes to remove the gelatin, treated with PBS1X, 1% triton, 2% serum for permeabilization and blocking. Primary antibodies were diluted in PBS1X 2% newborn calf serum and applied overnight at 4°C under a coverslip. Secondary antibodies were diluted in PBS1X and applied for 2 hours at room temperature. Washes were done in PBS1X. Antibodies used: custom-made polyclonal mouse anti-chick MMP14, mouse anti-AP2 (DSHB, 3B5), mouse anti-Snail2 (Cell Signaling, C19B7), rabbit anti-Sox10 (GeneTex, GTX128374), rabbit anti-PCM1 (Dammermann and Merdes, 2002), mouse anti-N-Cadherin (DSHB, 6B3), mouse anti-E-cadherin (BD Transduction Laboratories, 610181), mouse anti-phospho-histone 3 (Cell Signaling, MA312B), mouse anti-laminin (DSHB, 3H11).

### In situ hybridization

Wholemount in situ hybridization were performed as previously described (Theveneau 2007). Target sequence for the probe against chick-MMP14: 5’-cggcggcttcgataccatcgcggt Gctcaggggggagatgttcgtgttcaaggagcggtggctgtggcggctgcgggagcgccgggtgctgcccggttaccccctccctatggggcagctgtggcccggactgccccacagcatcgacgccgcctatgagaggaaggacggcaagttcgtcttcttcaaaggcgggcggcagtgggtgttctcggaggcggcgctgcagccgggcttcccgcgcgctctgccggacgtgggccgggggctgccggagcgcatcgacgccgcgctgctgtggctgcccagcggggccacgtacctcttccggggcgacaagtactaccggttcaatgaggagacggagtcggtggaccccgattaccccaaaagcatttccgtgtggggcggcgtccccgaatcaccccaaggagcatttatggggtcggatgacgcctacacgtactttgtgaagggctcccgctattggcagttcgacaaccgccagctgcgcgtcaccccgggttaccccaaatccctgctccgcgattgg-3’ corresponding to the portion of the coding sequence from 902-1454 (Yang et al., 2008) was amplified using forward primer: 5’ GACGGCGGCTTCGATACCA 3’ and reverse primer: 5’ ATCGCGGAGCAGGGATTT 3’. We also designed probes against portions 639-1052 (F: 5’ GTCACGACGTGTTCCTGGT 3’, R: 5’ GTCGATGCTGTGGGGCAGT 3’) and 196-565 (F: 5’ GGAAGTGTCGACCCGGAAA 3’, R: 5’ CGGGGAAGTAGGCGTGG 3’) which gave the same results. A probe against chick-Furin was generated using the following primers forward 5’ TCAGCTGGAGAGTAGTGGCT 3’, reverse 5’ CTCCTCAATGTCCGACTCCG 3’. Probes for Chicken TIMP2 and TIMP3 (Brauer and Cai, 2002), Foxd3 (Kos et al., 2001), Sox10 (Cheng et al., 2000), Ets1 (Vandenbunder et al., 1989), Integrin β3 (Pietri et al., 2003), Cadherin-6B and Cadherin-7 (Nakagawa and Takeichi, 1995).

### Chick MMP14 expression vectors

Full length MMP14 was synthetized using codon optimization to reduce the amount of GCs. Deletions and the point mutant forms were made by PCR strategies from the optimized sequence and subcloned in pCAGGS vector for electroporation.

### Image acquisition, treatment and analysis

Images were taken on a stereomicroscope Leica MZF10F equipped with a camera Leica DFC450C and the LAS software, a Nikon Eclipse 80i equipped with a camera DXM1200C and the NIS-elements software, a Zeiss AxioImager 2 equipped with a Hamamatsu ORCA Flash 4 camera and the Zen2 software, a confocal Zeiss 710 and a confocal Leica SP8. Images were then processed with FIJI.

### Statistics

Statistical analyses were performed with Prism 6 (GraphPad). Datasets were tested for Gaussian distribution. Student t-tests or ANOVA followed by multiple comparisons were used with the appropriate parameters depending on the Gaussian vs non-Gaussian characteristics of the data distribution. Significance threshold was set at p<0.05. Proportions were compared according to Taillard et al, 2008 (Taillard et al., 2008).

